# Fig4 confers tolerance to rapamycin in a Ste20-dependent manner independent of the Vac14 scaffold

**DOI:** 10.1101/2024.07.12.603206

**Authors:** Hannah E. Reeves, Anna King, Imran Khan, Asha Thomas, Corey Chung, Lauren D. Dotson, Anirudan Sivaprakash, Harrison A. Hall, Alim Habib, Sophia R. Lee, Bonnie C. Lloyd, Caroline L. Darbro, Bethany S. Strunk

## Abstract

**Summary Statement:** Disease-related Fig4 mutants displaying impaired association with the Fab1-Vac14-Fig4 complex confer tolerance to rapamycin in *Saccharomyces cerevisiae* independent of Vac14 in a Ste20-dependent manner. Fig4 can thus modulate homeostasis through association with pathways outside of the Fab1-Vac14-Fig4 complex in yeast.

The lipid phosphatase Fig4 modulates the production and turnover of the signaling lipid PI3,5P2 through participation in a complex with its opposing kinase, Fab1, and the scaffold protein Vac14. Fig4 point mutations are associated with neurological diseases for which there are currently no specific therapies. Pathology involving Fig4 is generally attributed to disruption of the Fab1-Vac14-Fig4 complex. Through genetic manipulations in yeast we show that expression of Fig4 disease-related mutants confers tolerance to rapamycin, an inhibitor of the master growth activator TORC1. This phenotype is dominantly conferred at high temperatures by Fig4 mutants that bind poorly to Vac14 or by excess wild-type Fig4. Rapamycin tolerance does not require Fig4 catalytic activity or Vac14, suggesting that Fig4 mediates this altered stress response through context-inappropriate binding interactions with unknown proteins. The p21-activated kinase Ste20 is specifically required for this novel Vac14-independent Fig4 function. Our study shows that Fig4 can modulate homeostasis in *Saccharomyces cerevisiae* through interactions outside the Fab1-Vac14-Fig4 complex.

## Introduction

Phosphoinositide (PI) lipid phosphatases modulate cellular signaling cascades through localized dephosphorylation of specific PI lipids on subcellular membrane domains. The PI phosphatase Fig4 (alpha **<u>F</u>**actor **<u>i</u>**nduced **<u>g</u>**ene **<u>4</u>**) (Erdman et al., 1998) is conserved in all eukaryotes and is important both for development and long-term homeostasis in mammals (Mironova etal., 2018; Lenk et al, 2016, 2011; Mironova et al, 2016; Chow et al, 2007). Homozygous loss of Fig4 in humans causes Yunis-Varon syndrome, a developmental disorder characterized by severe defects of the skeletal, nervous, and ectodermal tissue systems (Campeau et al, 2013; Nakajima et al, 2013; Umair et al, 2021). Biallelic mutations in Fig4 have been implicated in neurological disorders including Charcot-Marie-Tooth disease type 4J (CMT4J) (Chow et al, 2007), familial epilepsy with polymicrogyria (Baulac et al., 2014) and leukoencephalopathy (Lenk et al., 2019a; Sait et al., 2023). Single point mutations in Fig4 have been linked to amyotrophic lateral sclerosis (ALS) and primary lateral sclerosis (PLS) (Osmanovic et al., 2017; Chow et al., 2009; de Boer et al., 2023).

Fig4 dephosphorylates phosphatidylinositol 3,5-bisphosphate (PI3,5P2), thereby decreasing levels of this lipid by converting it to PI3P **(Figure 1A)** (Rudge et al., 2004; Lees et al., 2020). PI3,5P2 is a low abundance PI found primarily on late endosomes and lysosomes in mammalian cells and the vacuole in yeast (Rivero-Ríos and Weisman, 2022; Barlow-Busch et al., 2023). PI3,5P2 serves as an effector in diverse cellular pathways including control of actin dynamics, membrane homeostasis, and signaling in response to stress (Rivero-Ríos and Weisman, 2022). Effectors regulated through direct binding of PI3,5P2 include the general transcription factor Tup1, the lysosomal ion channels TRPML1 and TPC1/2, the vacuolar ATPase, and the Target of Rapamycin Complex 1 (TORC1) (McCartney et al., 2014).

**Figure 1:**
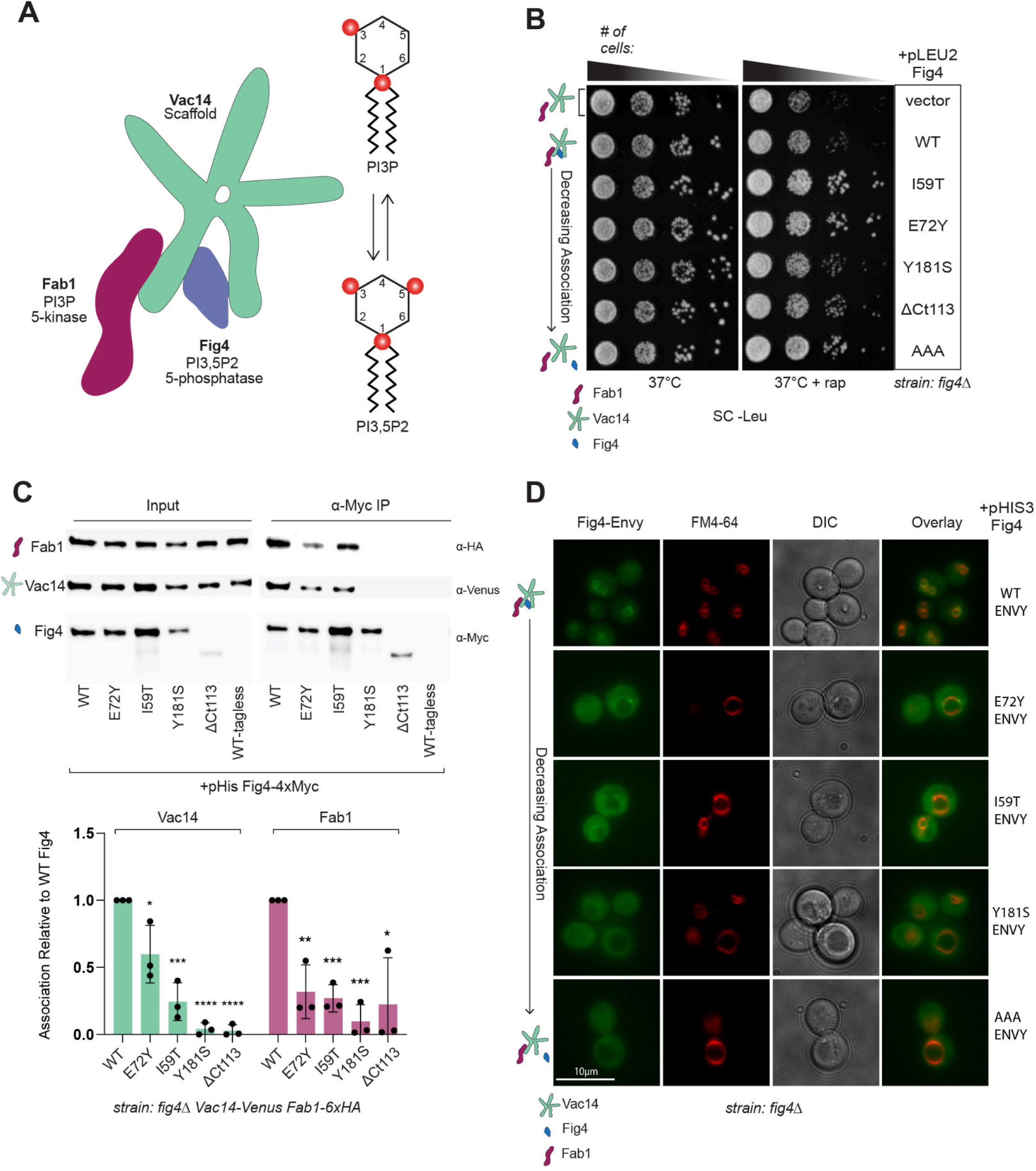
Fig4 mutants that have impaired association with Fab1 and Vac14 confer tolerance to rapamycin at 37°C. (A) Cartoon representation of Fab1-Vac14-Fig4 complex adapted from Lees 2020. Fig4 (∼100 kDa) and Fab1 (∼260 kDa) monomers associate with Vac14 pentamer (∼100 kDa x 5) to support the production and turnover of PI3,5P2 (Fig4 phosphatase activity removes the inositol 5-phosphate of PI3.5P2 to produce PI3P, Fab1 kinase activity performs the reverse reaction). (B) Disease-related Fig4 variants enhance rapamycin tolerance at 37°C. A *fig4Δ* strain was transformed with pRS415 (+pLEU2) plasmids expressing the indicated Fig4 variants with native promoter and 5′ and 3′ UTRs. Cells were spotted on SC-Leu agar plates in a 10-fold dilution series and incubated at 37°C for 2-3 days without rapamycin or 7-10 days with 10 nM rapamycin. (C) Fig4 disease-related variants display impaired association with Fab1 and Fig4. Western blot of proteins immunoprecipitated with anti-Myc antibody (IP) from a *fig4Δ Vac14-Venus Fab1-6xHA* strain transformed with pRS413 (+pHIS3) plasmids expressing indicated 4xMyc-tagged Fig4 variants or untagged Fig4 wild-type (WT-tagless) as a negative control. Plasmid based Fig4 expressed with native promoter and 5′ and 3′ UTRs. *Vac14-Venus* and *Fab1-6xHA* are expressed from endogenous loci. Bar graph shows quantification of association of Vac14 and Fab1 with Fig4 variants relative to Fig4-WT. Band intensities are normalized to Fig4 in each IP, with data points and mean from two independent experiments. Error bars represent standard deviation (ns not significant, *p<0.05, ** p < 0.01, *** p < 0.001, **** p < 0.0001 by two-tailed t-test) (D) Fig4 disease-related variants display diffuse cytoplasmic localization relative to Fig4-WT. Fluorescence micrographs of a fig4Δ strain transformed with pRS413 (+pHIS3) expressing C-terminally ENVY-tagged Fig4 variants with native promoter and 5′ and 3′ UTRs. Vacuoles were stained with FM4-64. Fig4 variants in this figure: Wild-type (WT), CMT4J (I59T), ALS (E72Y), and leukoencephalopathy (Y181S and ΔCt113), T52A-T62A-T78A (AAA) (Strunk et al., 2020), or no Fig4 (vector).

In addition to reducing PI3,5P2 levels, Fig4 promotes dynamic elevation of this substrate via participation in a protein complex harboring its opposing kinase Fab1. Fab1 is a PI 5-kinase **(Figure 1A)** (Sbrissa et al., 2007, 2008; Jin et al., 2008; Lees et al., 2020) and the sole enzyme catalyzing PI3,5P2 production in eukaryotic cells (Gary et al., 1998; Cooke et al., 1998). This complex is scaffolded on the Fab1 activator, Vac14. As a component of the Fab1-Vac14-Fig4 complex, Fig4 stimulates PI3,5P2 production independent of its catalytic activity and also likely via protein phosphatase function (Duex et al., 2006b; Lees et al., 2020). The molecular mechanisms by which Fig4 controls PI3,5P2 production and turnover are not fully understood but available data suggest that both processes require its stable and direct binding to the scaffold protein Vac14 (Rudge et al., 2004; Duex et al., 2006).

Fig4 null mutants are associated with PI3,5P2 insufficiency resulting in phenotypes that are often milder versions of those caused by loss of Fab1 or Vac14 (Lines et al., 2017; Duex et al., 2006b; Lenk et al., 2019b; Cao et al., 2023a; Chow et al., 2007). Accordingly, phenotypes associated with Fig4 mutations are generally attributed to dysregulation of PI3,5P2 due to impaired function of the Fab1-Vac14-Fig4 complex. Notably, overexpression of Fig4 can lead to different phenotypes than overexpression of Vac14 (Qi et al., 2022) and there are Fig4-dependent phenotypes that appear to be conferred independent of Fab1 and Vac14. For example, specific knockdown of Fig4, as opposed to Fab1 or Vac14, achieves growth arrest in triple negative breast cancer cell lines (Ikonomov et al., 2013). Moreover, knockdown of Fig4 in mouse adipocytes results in loss of insulin-dependent activation of the master growth regulator TORC1 without detectable changes in PI3,5P2 (Bridges et al., 2012). These studies raise questions as to the mechanism of Fig4 involvement in these specific phenomena. Although proteins uniquely associated with Fig4 have been reported (Qiu et al., 2021), potential roles for Fig4 outside of the Fab1-Vac14-Fig4 complex remain largely unexplored.

Here we report the unexpected finding that, independent of Vac14, disease-related mutants of Fig4 confer tolerance to the TORC1 inhibitor rapamycin in *Saccharomyces cerevisiae*. This phenotype is a dominant property of Fig4 mutants displaying impaired binding to the Fab1-Vac14-Fig4 complex and does not require Fig4 catalytic activity. We propose that Fig4-dependent tolerance to rapamycin results from functional, but context-inappropriate interactions between Fig4 and unrecognized protein partners when it is not tethered to Vac14. Rapamycin tolerance conferred by Fig4 requires the p21-activated kinase Ste20, a modulator of pathways controlling cytokinesis, morphogenesis, and signaling in response to multiple developmental and environmental stimuli (Boyce and Andrianopoulos, 2011; Chen and Thorner, 2007). Our findings show that Fig4 can modulate cellular homeostasis and proliferation in yeast independent of its known scaffold, Vac14.

## Results

### Disease-related variants of Fig4 confer tolerance to rapamycin at 37°C

Growth in S. *cerevisiae* is not limited by Fig4 function under a wide range of conditions (Giaever et al., 2002). When Fig4 is deleted from a wild-type laboratory strain, cells lacking Fig4 grow at the same rate as cells expressing wild-type Fig4 from a plasmid at 24°C or 37°C (Duex et al., 2006a; b; Strunk et al., 2020) **(Figure 1B)**. We were surprised to find that expression of several disease-related Fig4 variants enhanced growth relative to wild-type or no Fig4 when TORC1 was inhibited with 10 nM rapamycin at 37°C in this background. This Fig4-dependent tolerance to rapamycin was conferred by mutations associated with CMT4J, ALS, and PLS (I59T) (Chow et al, 2007; Mendes Ferreira et al, 2024; de Boer et al, 2023), ALS (E72Y) (Chow et al, 2009; Stump et al, 2023) and leukoencephalopathy (Y181S, ΔCt113) (Lenk et al, 2019a). A similar rapamycin tolerance was observed in cells expressing Fig4-T52AT62AT78A (Fig4-AAA) **(Figure 1B)**, a mutant not specifically linked to disease but where substitution mutations at three conserved threonines on the N-terminal surface of Fig4 near I59 and E72 disrupts binding of Fig4 to Vac14 (Strunk et al, 2020). Cells expressing no Fig4 displayed a growth rate similar to wild-type under these conditions. Therefore, rapamycin tolerance is not a result of loss of Fig4 function but is conferred by the presence of Fig4 mutants displaying behavior different from wild-type.

### Fig4 mutants conferring rapamycin tolerance are impaired in binding to the Fab1-Vac14-Fig4 complex

Fig4-E72Y and Fig4-I59T were shown previously to exhibit impaired association with the Fab1-Vac14-Fig4 complex as well as impaired stress-induced elevation of PI3,5P2 (Lenk et al, 2011; Chow et al, 2009). Association of Fig4-AAA with Fab1 and Vac14 is nearly undetectable and its impairment of stress-induced elevation of PI3,5P2 phenocopies Fig4 null mutants (Strunk et al, 2020). We sought to determine whether the other Fig4 variants conferring rapamycin tolerance are also impaired in association with Fab1 and Vac14. Western blots of cell lysates indicated that cellular levels of each Fig4-variant was similar; however, immunoprecipitation of Vac14 and Fab1 with the disease-related variants, I59T, E72Y, Y181S, and ΔCt113 was significantly impaired **(Figure 1C)**. Wild-type Fig4 localizes strongly to the outer membrane of the vacuole through its association with Vac14 (Rudge et al, 2004). To investigate whether disease-related variants of Fig4 were mis-localized, we performed fluorescence microscopy of live cells expressing Envy-tagged Fig4 variants. Fig4 mutants conferring rapamycin tolerance displayed weaker association with the vacuole and more diffuse cytoplasmic distribution relative to Fig4-WT reflecting their degree of impaired binding to Vac14 **(Figure 1D)**. As previously reported (Rudge et al, 2004; Duex et al, 2006a), cells expressing disease-related Fig4 variants displayed slightly enlarged vacuoles consistent with reduced PI3,5P2 levels in these cells **(Figure 1D)**.

### Fig4-dependent tolerance to rapamycin does not require Fig4 catalytic function

We recognized that rapamycin tolerance might be conferred through Fig4-mediated dephosphorylation of lipid or protein substrates, so we sought to determine whether rapamycin tolerance required Fig4 catalytic function. Substitution of the Fig4 catalytic cysteine with serine, Fig4-C467S, disrupts Fig4 catalytic function (Rohde et al., 2003; Lees et al., 2020) but does not prevent assembly of the Fab1-Vac14-Fig4 complex or stimulation of PI3,5P2 production (Strunk et al., 2020). To generate a catalytically-dead Fig4 mutant impaired in binding to Fab1 and Vac14, we introduced the C467S substitution into the Fig4-AAA variant (Fig4-AAA-C467S). Fig4-AAA-C467S displayed impaired binding to the Fab1-Vac14-Fig4 complex similar to that observed in Fig4-AAA **(Figure 2A)**. Fig4-AAA-C467S conferred rapamycin tolerance as well as Fig-AAA **(Figure 2B)** indicating that rapamycin tolerance is conferred independent of Fig4 catalytic function.

**Figure 2:**
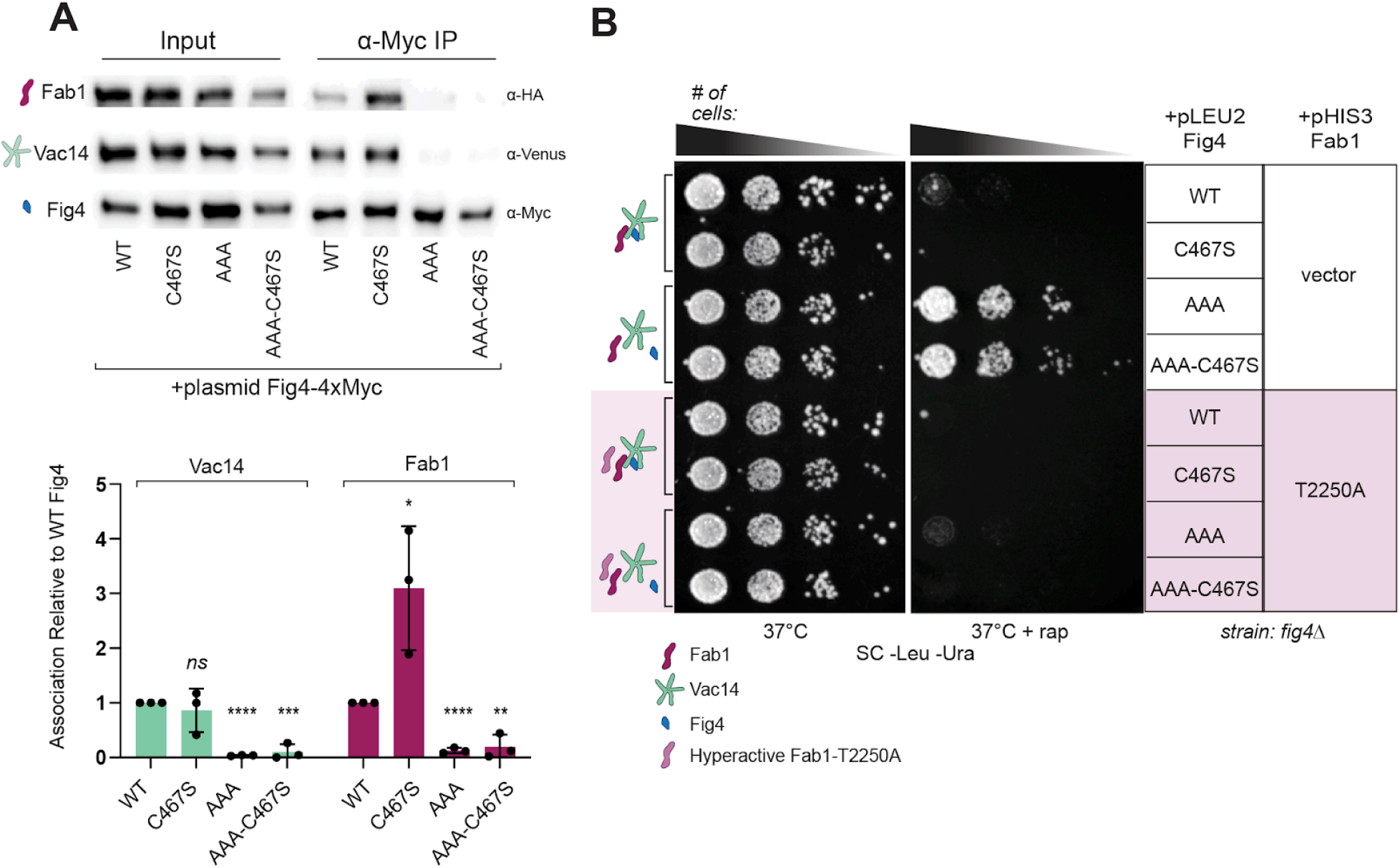
Fig4 catalytic function is not required for rapamycin tolerance at 37°C. **(A)** Catalytically-dead Fig4-C467S-T52A-T62A-T78A (AAA-C467S) is impaired in binding to Fab1 and Vac14. Western blot of proteins immunoprecipitated (IP) with anti-Myc antibody from a *fig4Δ Vac14-Venus Fab1-6xHA* strain transformed with pRS415 (+pLEU2) plasmids with native promoter and 5′ and 3′ UTRs expressing indicated 4xMyc-tagged Fig4 variants or untagged Fig4 wild-type (WT-tagless). Vac14-Venus and Fab1-6xHA are expressed from endogenous loci. The bar graph shows quantified association of Vac14 and Fab1 with Fig4 variants relative to Fig4-WT. Band intensities are normalized to Fig4-4xMyc in each IP, with data points and mean (bar) from three independent experiments. Error bars represent standard deviation *(ns* not significant, *p < 0.05, ** p < 0.01, *** p < 0.001, **** p < 0.0001 by two-tailed t-test). (B) Catalytically-dead Fig4 that binds poorly to Fab1 and Vac14 confers tolerance to rapamycin. A *fig4Δ* strain was co-transformed with pRS415 (+pLEU2) plasmids expressing the indicated Fig4 variants or no Fig4 (vector) and pRS416 (+pURA3) expressing hyperactive Fab1 (T2250A - pink shading) or no excess Fab1 in addition to wild-type Fab1 expressed from endogenous locus. Plasmid based Fig4 and Fab1 were expressed with native promoters and 5′ and 3′ UTRs. Cells were spotted on SC--Leu-Ura agar plates in a 10-fold dilution series and incubated at 37°C for 2-3 days without rapamycin or 7-10 days with 10 nM rapamycin. Fig4 variants in this figure: Wild-type (WT), catalytically-dead (C467S), T52AT62AT78A (AAA), catalytically dead C467S-T52AT62AT78A (AAA-C467S).

### Rapamycin tolerance conferred by Fig4 mutants is not a result of elevated PI3,5P2

Unexpectedly, we observed that relative to wild-type, expression of catalytically dead Fig4-C467S resulted in enhanced sensitivity to rapamycin in growth assays **(Figure 2B)**. Under standard growth conditions, cells lacking Fig4 have basal levels of PI3,5P2 similar to wild-type and are impaired in elevation of PI3,5P2 following hyperosmotic shock (Duex et al, 2006a). In contrast, expression of Fig4-C467S in yeast results in elevated basal levels of PI3,5P2 and sustained PI3,5P2 elevation following hyperosmotic stress (Strunk 2020). We hypothesized the enhanced rapamycin sensitivity observed in cells expressing Fig4-C467S could be a consequence of elevated PI3,5P2 production due to stable formation of the Fab1-Vac14-Fig4 complex in the absence of PI3,5P2-phosphatase activity. Co-expression of a hyperactive Fab1 mutant (Fab1-T2250A) displaying constitutive elevation of PI3,5P2 (Duex etal., 2006b; Lang et al., 2017) resulted in enhanced sensitivity to rapamycin at 37°C in the presence of all Fig4 variants tested **(Figure 2B)**. Therefore, it is unlikely that rapamycin tolerance conferred by Fig4 mutants is a result of elevation of PI3,5P2 in cells expressing Fig4 mutants.

### The rapamycin tolerance conferred by Fig4 mutants is dominant and is conferred by excess wild-type Fig4

Rapamycin tolerance persisted when we simultaneously expressed Fig4-WT and Fig4-AAA from separate centromeric plasmids, albeit to a lesser degree **(Figure 3A)**. The dominance of Fig4-AAA suggests that Fig4-AAA does not bind to the same binding sites in the Fab1-Vav14-Fig4 complex occupied by Fig4-WT. Otherwise, simultaneous expression of these variants would be expected to result in a wild-type phenotype because Fig4-WT would outcompete Fig4-AAA. We therefore considered the possibility that rapamycin tolerance is conferred by elevated levels of free Fig4 in mutants impaired in association with Fab1 and Vac14. In support of this model, rapamycin tolerance was recapitulated by expressing Fig4-WT from two centromeric plasmids, modestly increasing the cellular level of Fig4 protein **(Figure 3A, Figure S1)**. Taken together, these data imply that the ability to confer rapamycin tolerance is not unique to Fig4 mutants but rather a general property of Fig4 protein not engaged with the Fab1-Vac14-Fig4 complex, either as a result of mutation or when present in excess.

**Figure 3:**
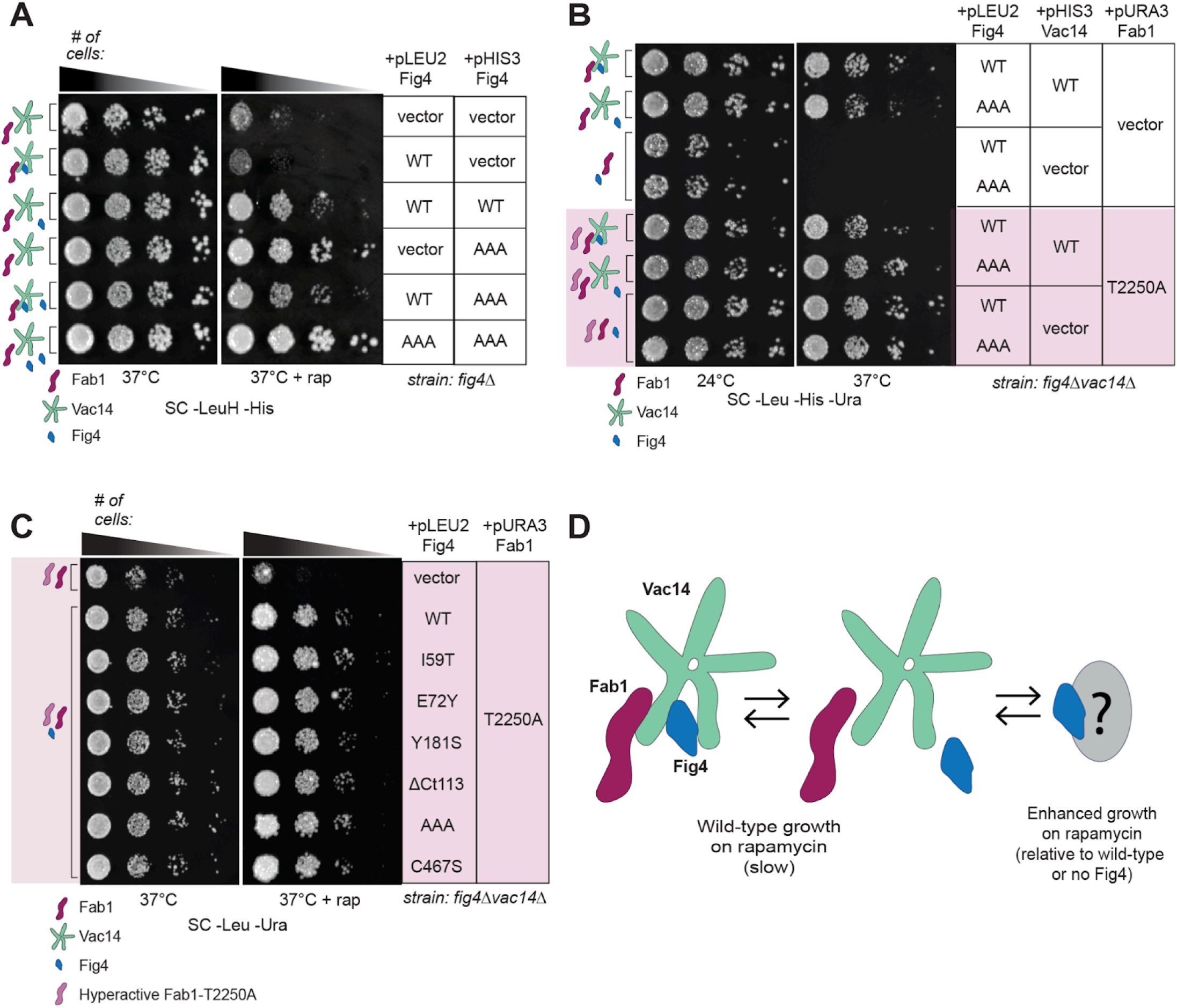
Fig4 variants confer rapamycin tolerance independent of Vac14. (A) The growth advantage conferred by Fig4 variants is dominant to wild-type Fig4 and is also conferred by wild-type Fig4 when in excess. A fig4Δ strain was co-transformed with pRS415 (+pLEU2) and pRS413 (+pHIS3) plasmids expressing the indicated Fig4 variants or no Fig4 (vector) with native promoter and 5′ and 3′ UTRs. Cells were spotted on SC-Leu-His agar plates in a 10-fold dilution series and incubated at 37°C for 2-3 days without rapamycin or 7-10 days with 10 nM rapamycin. (B) Temperature sensitivity of a *fig4Δvac14Δ* strain is rescued by expression of hyperactive Fab1-T2250A. A *fig4Δvac14Δ* strain was co-transformed with pRS415 (+pLEU2) plasmids expressing the indicated Fig4 variants or no Fig4 (vector), pRS413 (+pHIS3) plasmids expressing the wild-type Vac14 or no Vac14 (vector), and pRS416 (+pURA3) expressing hyperactive Fab1 (T2250A - pink shading), or no Fab1, in addition to wild-type Fab1 expressed from endogenous locus. Plasmid based Fig4, Vac14, and Fab1 were expressed with native promoters and 5′ and 3′ UTRs. Cells were spotted on SC-Leu-His-Ura agar plates in a 10-fold dilution series at 24°C and 37°C. (C) Fig4 variants, including wild-type and catalytically-dead Fig4, confer tolerance to rapamycin in the absence of Vac14. A *fig4Δvac14Δ* strain was co-transformed with pRS415 (+pLEU2) plasmids expressing the indicated Fig4 variants or no Fig4 (vector) and pRS416 (+pURA3) expressing hyperactive Fab1 (T2250A) in addition to wild-type Fab1 expressed from endogenous locus. Plasmid based Fig4 and Fab1 were expressed with native promoters and 5′ and 3′ UTRs. Cells were spotted on Sc-Leu-Ura agar plates in a 10-fold dilution series and incubated at 37°C for 2-3 days without rapamycin or 7-10 days with 10 nM rapamycin. (D) Model for Fig4-dependent tolerance to rapamycin. When Fig4 is not bound to the Fab1-Vac14-Fig4 complex through direct binding with Vac14, it is available to physically associate with molecular components of another cellular pathway resulting in enhanced growth on 10 nM rapamycin at 37°C. Fig4 variants in this figure: Wild-type (WT), CMT4J (I59T), ALS (E72Y), and leukoencephalopathy (Y181S and ΔCt113), T52A-T62A-T78A (AAA), catalytically-dead (C467S).

### Fig4 confers rapamycin tolerance in the absence of Vac14

Available data suggest that association between Fig4 and Fab1 requires direct binding between Fig4 and the scaffold protein Vac14 (Jin et al., 2008; Strunk et al., 2020). If rapamycin tolerance is conferred by Fig4 not engaged with the Fab1-Vac14-Fig4 complex, any Fig4 variant would be expected to confer rapamycin tolerance in the absence of Vac14. Testing this prediction is complicated by the fact that cells lacking Vac14 do not grow at 37°C due to PI3,5P2 insufficiency (Dove et al., 2002; Bonangelino et al., 1997; Murén et al., 2001; Gomes de Mesquita et al., 1996). To rescue growth in a *vac14Δfig4Δ* strain at 37°C, we expressed the Fab1-T2250A mutant to restore sufficient PI3,5P2 production in these cells (Duex et al., 2006b) **(Figure 3B)**. In this rescued background lacking Vac14, all Fig4 variants tested, including Fig4-WT and Fig4-C467S, were more tolerant to rapamycin at 37°C than cells expressing no Fig4 **(Figure 3C)**. While there is no reported evidence for Fig4 associating with Fab1 in the absence of Vac14, we cannot rule out the possibility that Fig4 confers rapamycin tolerance through an uncharacterized interaction with Fab1. Regardless, the data demonstrate that Fig4 enhances growth on rapamycin independent of Vac14 and its canonical association with the Fab1-Vac14-Fig4 complex.

### The p21-activated protein kinase Ste20 is required for rapamycin tolerance conferred by Fig4 mutants

We hypothesize that Fig4 disease-related mutants interact with a previously unrecognized protein partner or partners in a context-inappropriate manner to confer rapamycin tolerance **(Figure 3D)**. The p21-activated protein kinase Ste20 was among the candidates for such a protein because of its involvement in two Fig4-related pathways: response to mating pheromone (Elion, 2000) and acclamation to hyperosmotic stress (Raitt et al., 2000). Moreover, a genetic interaction between Fig4 and Hippo, the Ste20 homolog in flies, was recently reported (Kushimura et al., 2018). Fig4-dependent rapamycin tolerance is ablated in cells deleted for Ste20 indicating that it is indeed required for this phenotype **(Figure 4A)**. Importantly, Ste20 is also required for rapamycin tolerance conferred by Fig4 in the absence of Vac14 **(Figure 4B)** supporting the likelihood that enhanced growth is conferred by Fig4 through the same mechanism both in the presence and absence of Vac14. Ste20 has a close homolog, Cla4, with which it shares some redundancy such that the double knockout of Ste20 and Cla4 is lethal (Keniry and Sprague, Jr., 2003; Cvrcková et al., 1995; Bartholomew and Hardy, 2009; Yau et al., 2017). Although a *fig4Δcla4Δ* double knockout grew slowly, enhanced growth on rapamycin at 37°C in cells expressing Fig4-AAA persisted **(Figure S2)** suggesting that support of rapamycin tolerance is a role unique to Ste20. To investigate the possibility that Ste20 mediates rapamycin tolerance through a physical interaction with Fig4, we attempted to co-immunoprecipitate Ste20 with Fig4 but were unable to detect an interaction (data not shown).

**Figure 4:**
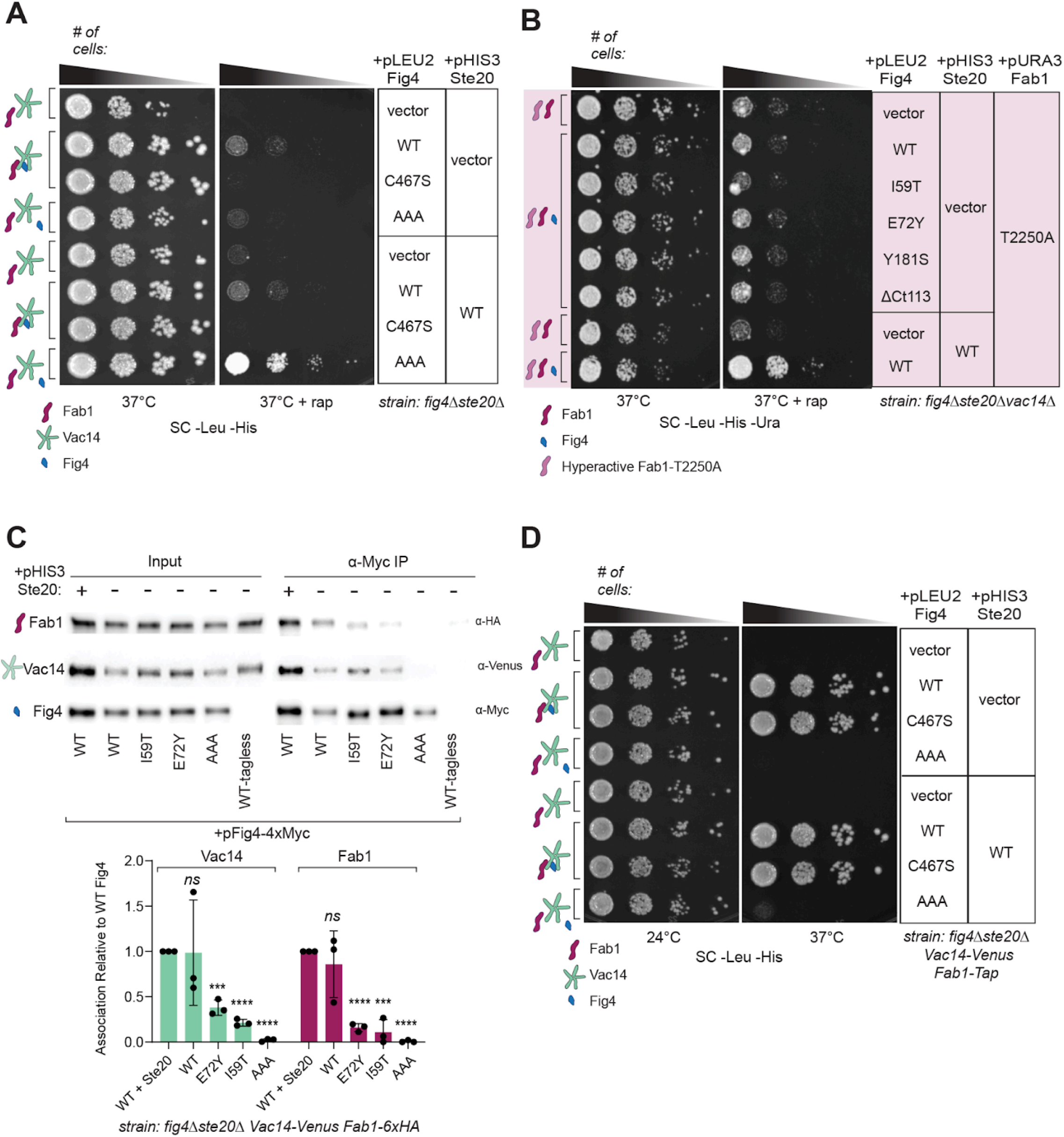
Ste20-kinase activity is required for Fig4-dependent tolerance to rapamycin but not for Fab1-Vac14-Fig4 related functions of Fig4. **(A)** Ste20 is required for the tolerance to rapamycin conferred by Fig4. A *fig4Δste20Δ* strain was co-transformed with pRS415 (+pLEU2) plasmids expressing the indicated Fig4 variants or no Fig4 (vector) and pRS413 (+pHIS3) expressing wild-type Ste20 (WT) or no Ste20 (vector). Plasmid based Fig4 and Ste20 were expressed with native promoters and 5′ and 3′ UTRs. Cells were spotted on SC-Leu-His agar plates in a 10-fold dilution series and incubated at 37°C for 2-3 days without rapamycin or 7-10 days with 10 nM rapamycin. (B) Ste20 is required for Fig4-dependent tolerance to rapamycin in the absence of Vac14. A *fig4Δste20Δvac14Δ* strain was co-transformed with pRS415 (+pLEU2) plasmids expressing the indicated Fig4 variants or no Fig4 (vector), pRS413 (+pHIS3) plasmids expressing wild-type Ste20 (WT) or no Ste20 (vector), and a pRS416 (+pURA3) expressing hyperactive Fab1 (T2250A - pink shading). Plasmid based Fig4, Ste20, and Fab1 were expressed with native promoters and 5′ and 3′ UTRs. Cells were spotted on SC-Leu-His-Ura agar plates in a 10-fold dilution series and incubated at 37°C for 2-3 days without rapamycin or 7-10 days with 10 nM rapamycin. (C) Ste20 is not required for Fig4 association with Fab1 and Vac14. Western blot of proteins immunoprecipitated with anti-Myc antibody from a *fig4Δste20Δ* strain co-transformed with pRS415 (+pLEU2) plasmids expressing the indicated Fig4 variants or no Fig4 (vector) and pRS413 (+pHIS3) expressing wild-type Ste20 (WT) or no Ste20 (vector). Plasmid based Fig4 and Ste20 were expressed with native promoters and 5′ and 3′ UTRs. Vac14-Venus and Fab1-6xHA are expressed from endogenous loci. Bar graph shows quantification of association of Vac14 and Fab1 with Fig4 variants relative to Fig4-WT. Band intensities are normalized to Fig4 in each IP, with data points and mean from three independent experiments. Error bars represent standard deviation (ns not significant, *** p < 0.001, **** p < 0.0001 by two-tailed t-test). (D) Ste20 is not required for Fig4-dependent growth in a strain background where Fig4 association with the Fab1-Vac14-Fig4 complex is required at 37°C (Vac14-Venus Fab1-TAP, Strunk et al., 2020). A *fig4Δste20Δ Vac14-Venus Fab1-TAP* strain was co-transformed with pRS415 (+pLEU2) plasmids expressing the indicated Fig4 variants or no Fig4 (vector) and pRS413 (+pHIS3) expressing wild-type Ste20 (WT) or no Ste20 (vector). Vac14-Venus and Fab1-TAP were expressed from endogenous loci. Growth assays performed as in panel A.

### Ste20 is not required for Fab1-Vac14-Fig4 complex associated functions of Fig4

We next investigated the possibility that the loss of Fig4-dependent rapamycin tolerance in the absence of Ste20 was due to a generalized loss of Fig4 function in this background. Western blots of cell lysates lacking Ste20 indicated that Fig4, Vac14, and Fab1 are stably expressed **(Figure 4C)**. Moreover, immunoprecipitations of Fab1 and Vac14 with Fig4 from these cells indicated that Fab1 and Vac14 are efficiently pulled down with Fig4-WT both in the presence and absence of Ste20 **(Figure 4C)**. Rapamycin tolerance conferred by Fig4 mutants showed reduced association with Fab1 and Vac14 relative to Fig4-WT in the absence of Ste20 **(Figure 4C)**, similar to what was observed in its presence (see Figure 1). We previously published growth assays in a yeast strain where growth at 37°C is dependent on Fig4 as a result of C-terminal tags on genomic copies of Fab1 and Vac14 *(fig4Δ Fab1-Tap Vac14-Venus)* (Strunk et al., 2020). In this strain background, the ability of Fig4 variants to support growth is directly correlated with the strength of their association with Fab1 and Vac14 and their ability to support production of PI3,5P2 following hyperosmotic shock (Strunk et al., 2020). Deletion of Ste20 does not alter the ability of Fig4 variants to support growth in this background **(Figure 4D)**. Taken together, these data suggest Ste20 is specifically required for Fig4 dependent rapamycin tolerance and not for Fig4 functions involving the Fab1-Vac14-Fig4 complex.

## Discussion

The data presented here show that Fig4 can influence cellular homeostasis in S. *cerevisiae* independent of its known scaffold protein, Vac14. Fig4 confers enhanced tolerance to rapamycin at 37°C when it is dissociated from the Fab1-Vac14-Fig4 complex. This phenotype is not due to failure to turn over PI3,5P2 nor any other Fig4-phosphatase function. Moreover, it is not likely a result of general elevation of PI3,5P2 as genotypes associated with elevated PI3,5P2 display enhanced rapamycin sensitivity. Instead, our data strongly suggest that Fig4 not engaged with the Fab1-Vac14-Fig4 complex is available to interact with an alternative cellular pathway in a context-inappropriate manner to promote growth under these conditions. The requirement for Ste20 for this Fig4-dependent phenotype suggests that Fig4 modulates this phenotype through a physical interaction with components of a Ste20-dependent pathway.

We attempted to detect, but were unable to confirm, a physical interaction between Ste20 and Fig4. We performed immunoprecipitations with cells grown under a number of conditions in liquid culture. It is possible that these conditions could not recapitulate the cellular context of colonies on agar plates rapamycin at 37°C. Moreover, membrane-associated interactions may have been missed in the lysis conditions used. It is worth noting that the experiments we performed to detect interactions of Fig4 with Fab1 and Vac14 had similar limitations. However, the persistence of Fig4-dependent tolerance to rapamycin in the absence of Vac14 supports a model where Fig4 that is not bound to Vac14 can confer this phenotype.

An additional limitation to our study is that we did not measure PI3,5P2. Although tolerance to rapamycin is conferred by Fig4 independent of Vac14, we cannot rule out that it involves Fab1 and regulation of PI3,5P2 production. Notably, it was recently reported that TORC1-dependent phosphorylation of Fab1 shifts its localization from the vacuole membrane to vacuole-adjacent signaling endosomes (Chen et al., 2020). Restricting Fab1 to one compartment or the other results in divergent responses to rapamycin at 37°C (Chen et al., 2020) similar to what we report here. If general reduction or elevation of PI3,5P2 is not responsible for Fig4-dependent rapamycin tolerance, it is possible that Fig4 modulates Fab1 function in specific contexts or compartments independent of Vac14. Notably, the existence of independently regulated pools of PI3,5P2 with distinct effectors was proposed to explain how Fig4 knockdown led to loss of TORC1 activation in mouse adipocytes without detectable changes in global PI3,5P2 levels (Bridges et al., 2012). Fig4 and Ste20 could play a role in tuning PI3,5P2 production in these distinct compartments.

This is not the first study linking Fig4 and Ste20. A genetic interaction between Fig4 and the *Drosophila* Ste20 homolog Hippo was previously reported (Kushimura et al., 2018). Ste20 functions as a MAP4K in pathways including mating, osmoregulation, cell wall integrity, and filamentous growth in response to stimuli at the plasma membrane (Boyce and Andrianopoulos, 2011; Chen and Thorner, 2007). Fig4 is specifically associated with mating and hyperosmotic shock in yeast (Erdman et al., 1998; Duex et al., 2006a). Ste20 also functions in vacuole inheritance, mitotic exit, modulation of the vacuolar ATPase, programmed cell death, and regulation of sterol uptake (Lin et al., 2009; Du and Liang, 2006; Ahn et al., 2005; Lin et al., 2012; Bartholomew and Hardy, 2009; Yau et al., 2017; Durant et al., 2024). Although Fig4 has not be directly implicated in these pathways, PI3,5P2 and TORC1 have been associated with several (Huda et al., 2023; Alhaj Sulaiman et al., 2022; Rivero-Rios and Weisman, 2022; Worley et al., 2015; Li et al., 2014; Okreglak et al., 2023; Wible et al., 2024; Groth et al., 2022).

Human neuropathies related to mutations in Fig4 involve demyelination and axon loss in the peripheral or central nervous systems (Hu et al., 2018; Lenk et al., 2019a; Chow et al., 2009). Differentiation of oligodendrocytes and Schwann cells during development and the repair of the myelin sheath in mammals has been shown previously to involve each of the pathways implicated here: Fig4-dependent production and turnover of PI3,5P2, Ste20-related kinases, and the switch between catabolic and anabolic processes through TORC1 (Cherchi et al., 2021; Figlia et al., 2018, 2017; Hu et al., 2016; Brown et al., 2021). Our data suggest that signals from these pathways converge on a novel Vac14-independent function of Fig4 in S. *cerevisiae*. Rapamycin tolerance is conferred in yeast at human body temperature by Fig4 mutants associated with human disease. It remains to be determined whether this Vac14-independent function of Fig4 is conserved in humans. If such interactions exist, they could contribute to disease-pathology caused by point mutations in Fig4.

In contrast to what we observed in yeast, Fig4 mutants impaired in binding to Vac14 are efficiently degraded in mammalian cells (Lenk et al., 2011; Ikonomov et al., 2010). Pathology is accordingly attributed to loss of Fig4 function and disruption of the Fab1-Vac14-Fig4 (PIKfyve-ArPIKfyve-Sac3) complex. Nonetheless, a Fab1-independent role for Fig4 in maintaining protein levels of specific surface expressed glycoproteins in mouse macrophage cells was previously reported (Morioka et al., 2017). Moreover, upregulation of Fig4 in triple negative breast cancer drives proliferation in cell lines without significant effects on PI3,5P2 levels (Ikonomov et al., 2013). Neither Fab1 nor Vac14 knockdown can achieve the growth arrest mediated by direct knockdown of Fig4 in these cells (Ikonomov et al., 2013) suggesting that Fig4 independently drives proliferation. It is possible that when Fig4 levels are sufficiently high there is enough free Fig4 to influence growth through interactions with a Vac14-independent pathway. A recent study showed that overexpression of catalytically-dead or wild-type Fig4, but not Vac14, leads to delocalization of the neuron specific PI3,5P2 effector NSG1 in rodent neuronal cell lines (Qi et al., 2022). It is not known whether this reflects increased binding of Fig4 to Vac14 or Vac14-independent interactions of Fig4 in these cells. It is notable that autosomal dominant mutations in Fig4 have been associated with ALS (Yilihamu et al., 2022; Chow et al., 2009; Ko et al., 2020) raising the possibility that disease may not simply be a result of loss of Fig4 function in these cases.

TORC1 is a master growth regulator activated under favorable growth conditions to promote anabolic processes such as growth and cell proliferation while inhibiting catabolic processes including autophagy and cell death (Khalil et al., 2023; Foltman and Sanchez-Diaz, 2023). Rapamycin-mediated inhibition of TORC1 induces a general stress response including changes in gene expression that mimic nutrient starvation (Alfatah et al., 2021; Hughes Hallett et al., 2014; Bandhakavi et al., 2008). Wild-type yeast cells have evolved to grow slowly under these conditions. Rapamycin tolerance therefore may reflect maladaptive signaling, resulting in enhanced growth under conditions where slow growth increases likelihood of survival. Given that wild-type Fig4 supports rapamycin tolerance when it is not associated with Vac14, we hypothesize that there are developmental and environmental contexts in which the regulated dissociation of Fig4 from Fab1 and/or Vac14 facilitates adaptive signaling to maintain cellular homeostasis. The aims of future studies will include identification of conditions under which Fig4 dissociates from Vac14 and the mechanistic basis for this dissociation.

## Materials and Methods

### Yeast strains and plasmids

Strains and plasmids used in this study are listed in Table 1. All strains come from the same haploid yeast background, LSWY3250 (derived from SEY6210). All plasmids are centromeric (Sikorski and Hieter, 1989). Target gene deletions generated for this study were made by PCR generation of a homology cassette using a KanMX6 (STE20) or a NatMX6 (CLA4) resistance module as a dominant marker (Goldstein and McCusker, 1999; Longtine et al., 1998). Yeast were grown in yeast extract-peptone-dextrose (1% yeast extract, 2% peptone, 2% dextrose; YEPD) or synthetic complete (SC) minimal medium without selective amino acids at the indicated strains then transformed with plasmids expressing variants of Fig4, Vac14, Ste20 and/or Fab1 from their native promoters and 5’ and 3’UTRs. Transformed cells were grown overnight in selective SC media to mid-log phase. Cultures were diluted to ∼1.5 × 10^6^ cells per mL a 10-fold serial dilution was plated on selective SC agar plates (1.5%) using a 48-Pin Microplate Replicator (V & P Scientific, Inc). Plates were grown at 24°C or 37°C for 3 days without rapamycin or 7-10 days with 10 nM rapamycin.

**Table 1.**
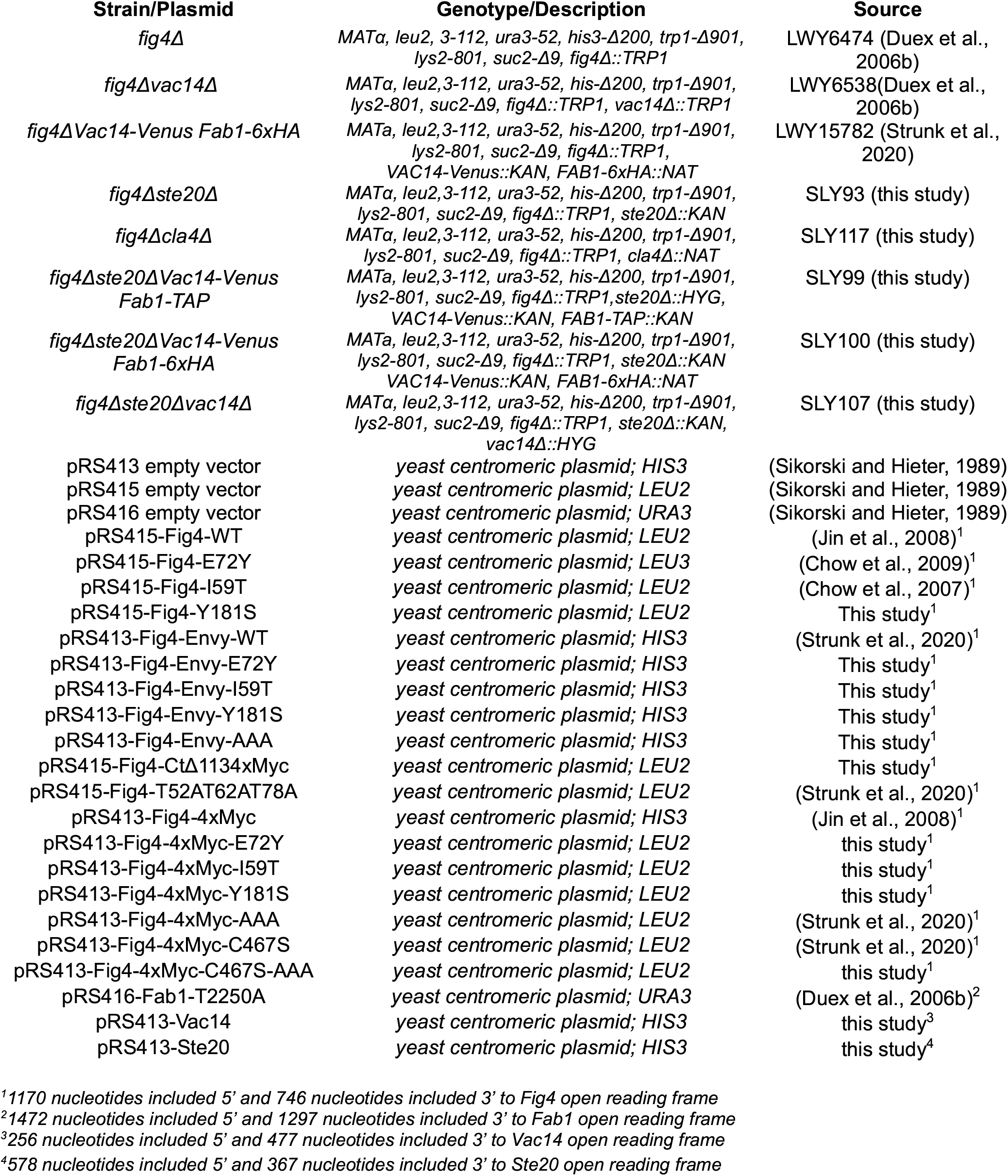
Strains and plasmids used in this study.

### Fluorescence microscopy

Yeast were grown in selective SC medium to an OD600 of 0.5 at 24°C and labeled with FM4-64 (Vida and Emr, 1995). Cells were harvested at 1000 × g for three minutes at 24°C and resuspended in 250 μl YEPD. 6μl FM4-64 (BIOTIUM) at 2mg/mL in DMSO was added and cells were incubated with shaking at 24°C for 30 minutes. FM4-64-labeled cells were washed twice with 250μl SC media then resuspended in 4 mL room temperature selective SC media. 1 mL cells were sedimented in microfuge tubes at 1000 × g for three minutes and gently resuspended in 20-50 μl media. 2-3μl of cells were applied to glass micro slides (Leica 3800220) imaged under cover glass (Leica 3800101) with a Nikon Eclipse TE2000-U Inverted Phase Contrast DIC fluorescent microscope using a Nikon Plan Apo VC 100x Oil Immersion Microscope Objective (n.a. 1.4 - Nikon MRD01901) with Cargille Laser Liquid 5610 (Cargille Labs 20130,20127-RCF). Images were captured with a Photometries CoolSNAP HQ Monochrome camera using NIS-Elements software interface.

### Co-immunoprecipitations from yeast

Yeast strains (fig4ΔVac14-VenusFab1-6xHA or fig4Δste20ΔVac14-VenusFab1-6xHA) were transformed with plasmids expressing variants of 4xMyc-tagged Fig4 or tagless Fig4 and additional plasmids as indicated. Log-phase cells (25 OD600 U) grown in 50 mL liquid culture at 24°C were harvested and lysed with ½ volume 0.5 mm diameter zirconia/silica beads (Biospec; 11079105z) and 3 volumes (3 μL per mg of cells) of IP buffer (50 mM Tris, pH 7.5, 120 mM NaCI, 10 mM EDTA, 1 mM EGTA, 5 mM 2-glycerophosphate (Sigma; G9422), 1x complete protease inhibitor cocktail EDTA-free (Roche; 11836170001), and 1x Protease Inhibitor Cocktail for use with fungal and yeast extracts (Sigma; P8215), 3 mM benzamidine, 1 μg/ml leupeptin, 2 μg/ml aprotinin, and 6 μg/ml chymostatin, 1 ug/mL pepstatin). All subsequent steps were carried out at 4°C. Cells were disrupted for 8 × 1 min with a SoniBeast - Small Sample Cell Disruptor (Biospec; 42105) with 2 min intervals in an ice water bath. Debris was removed by centrifugation for 5 min at 500 × g. The supernatant was mixed with 5% octyl-glucoside (Sigma; 08001) in lysis buffer for a final concentration of 0.5% octyl-glucoside and incubated for 30 min. Octyl-glucoside-solubilized lysate was cleared by spinning at 13,000 × g for 10 min. Supernatants were incubated with 0.8 pl per 100 pl lysate mouse anti-Myc antibody clone 9E10 (Sigma; 05-419) for 1 hour. 100μl of each lysate was applied to 15 μl Protein-G Sepharose beads (Cytiva; 17061801) equilibrated in lysis buffer and incubated with rocking for 1 h. Beads were washed three times with 500 μl of IP buffer containing 0.5% octyl-glucoside, 1 mM benzamidine, 5 mM 2-glycerophosphate, and 1 × Roche Complete inhibitor cocktail. Bound protein was eluted by heating immunoglobulin G beads with 35 μl of denaturing urea SDS-loading dye (1% SDS, 8 M urea, 10 mM Tris, pH 6.8, 10 mM EDTA, 0.01% bromophenol blue, 1x Roche Complete inhibitor cocktail, 1 mM benzamidine, 1 μg/ml leupeptin, 2 μg/ml aprotinin, and 6μ g/ml chymostatin, 1 ug/mL pepstatin and 5% 2-Mercaptoethanol) at 85°C for 10 min and spun for 1 min at 1000 × g prior to SDS-PAGE and Western blot analysis.

### Quantification of relative levels of Fig4 protein in cells

Yeast were grown in selective SC medium to an OD600 of 0.5 at 24°C. 1 mL cells were harvested at 1000 × g for 3 min at room temperature. Cells were lysed with 100 μl denaturing urea SDS-loading dye (1% SDS, 8 M urea, 10 mM Tris, pH 6.8, 10 mM EDTA, 0.01% bromophenol blue, 1x complete protease inhibitor cocktail EDTA-free (Roche; 11836170001), 1 mM benzamidine, 1 μg/ml leupeptin, 2 μg/ml aprotinin, and 6 μg/ml chymostatin, 1 μg/mL pepstatin and 5% 2-Mercaptoethanol). 50 μL of 0.5 mm diameter zirconia/silica beads (Biospec; 11079105z) were added to each sample before cells were disrupted with a SoniBeast - Small Sample Cell Disruptor (Biospec; 42105) in 0.5 ml tubes for 10 minutes at 4°C. Debris was removed by centrifugation for 5 min at 500 × g. Tubes were immediately heated at 95°C for 10 min and spun for 1 min at 1000 × g. prior to SDS-PAGE and Western blot analysis.

### Western Blot Analysis

Each sample was run on a 4-20% SDS-PAGE gel (BioRad; 4561096) and transferred on to nitrocellulose membranes (Cytiva; 10600096). The following primary antibodies were used to detected affinity tagged constructs: Fig4-4xMyc with rabbit anti-Myc at 1:1000 (Cell Signaling; 2278S), Fab1-6xHA with rabbit anti-HA at 1:1000 dilution in 5% nonfat dry milk in TBST. (Cell Signaling; 3724S), and Vac14-Venus with mouse anti-GFP 1:1000 dilution in 5% nonfat dry milk in TBST (Sigma; 11814460001). Detection through chemiluminescence was implemented with horseradish peroxidase conjugated secondary antibodies (Jackson Immunoresearch Laboratories; donkey anti-mouse IgG - 102650-014 or goat anti-rabbit IgG - 102645-188 at 1:10,000 dilution in 5% nonfat dry milk in TBST) and ECL Prime Western Blotting Detection Reagents (Amersham; RPN2232). Chemiluminescent blots were imaged with a Licor Odyssey Fc Imager using LICORbio Image Studio Software. For Co-immunoprecipitations, Fab1 and Vac14 bands were normalized to Fig4 in the corresponding immunoprecipitation. Association is expressed relative to Fig4-Myc wild-type in each experiment. For relative quantification of Fig4-4xMyc in lysates, bands were normalized to PGK1 detected with a mouse anti-Pgk1 antibody at 1:10,000 dilution in 5% nonfat dry milk in TBST (Invitrogen; 459250). Fig4 levels are expressed relative to wild-type Fig4 expressed from a single centromeric plasmid (1 copy wild-type).

### Statistical analysis

All statistical analysis was done using GraphPad Prism. Statistical comparison between Fig4 variants and wild-type (for immunoprecipitations) or one copy Fig4-WT (for determination of relative levels of one or two copies of Fig4 variants) was conducted using the two-sided Student’s t test, as indicated in the figure legends. Differences were considered significant if the P value was <0.05 *(ns* not significant, *p < 0.05, ** p < 0.01, *** p < 0.001, **** p < 0.0001).

### Online supplemental material

Fig. S1 shows the relative protein levels of Fig4 expressed from one or two centromeric plasmids to accompany Figure 3. Fig. S2 shows the persistence of relative tolerance to rapamycin conferred by Fig4-AAA relative to Fig4-WT and no Fig4 in the absence of the Ste20 homolog Cla4.

## Acknowledgements

This research was funded by the Voelcker Fund Young Investigator Award (MMTVF20214) and an NIH - NIGMS R00 award (GM120511-03) We thank Dr. Lois Weisman for her mentorship and inspiration for this project. We thank Dr. Katrin Karbstein, Dr, Jason A. MacGurn, Dr. Nathaniel Hepowit, Dr. Luis Giavedoni, Dr. Benjamin Tu, and Cole McGuire for advice and feedback, and Dr. Lois Weisman and Dr. Benjamin Tu for strains and plasmids.

## Author contributions

H.E.R.: investigation & conceptualization, A.K.: investigation, visualization, writing - review & editing, & formal analysis, LK.: investigation & conceptualization, A.T.: investigation, C.C.: investigation, A.S.: investigation, visualization & writing - review & editing, L.D.D.: investigation & visualization, H.A.H: investigation, A.H.: investigation, S.R.L.: investigation, B.C.L.: investigation, C.L.D.: investigation, B.S.S.: conceptualization, investigation, formal analysis, and writing - original draft

## Supplementary Material

**Figure S1.**
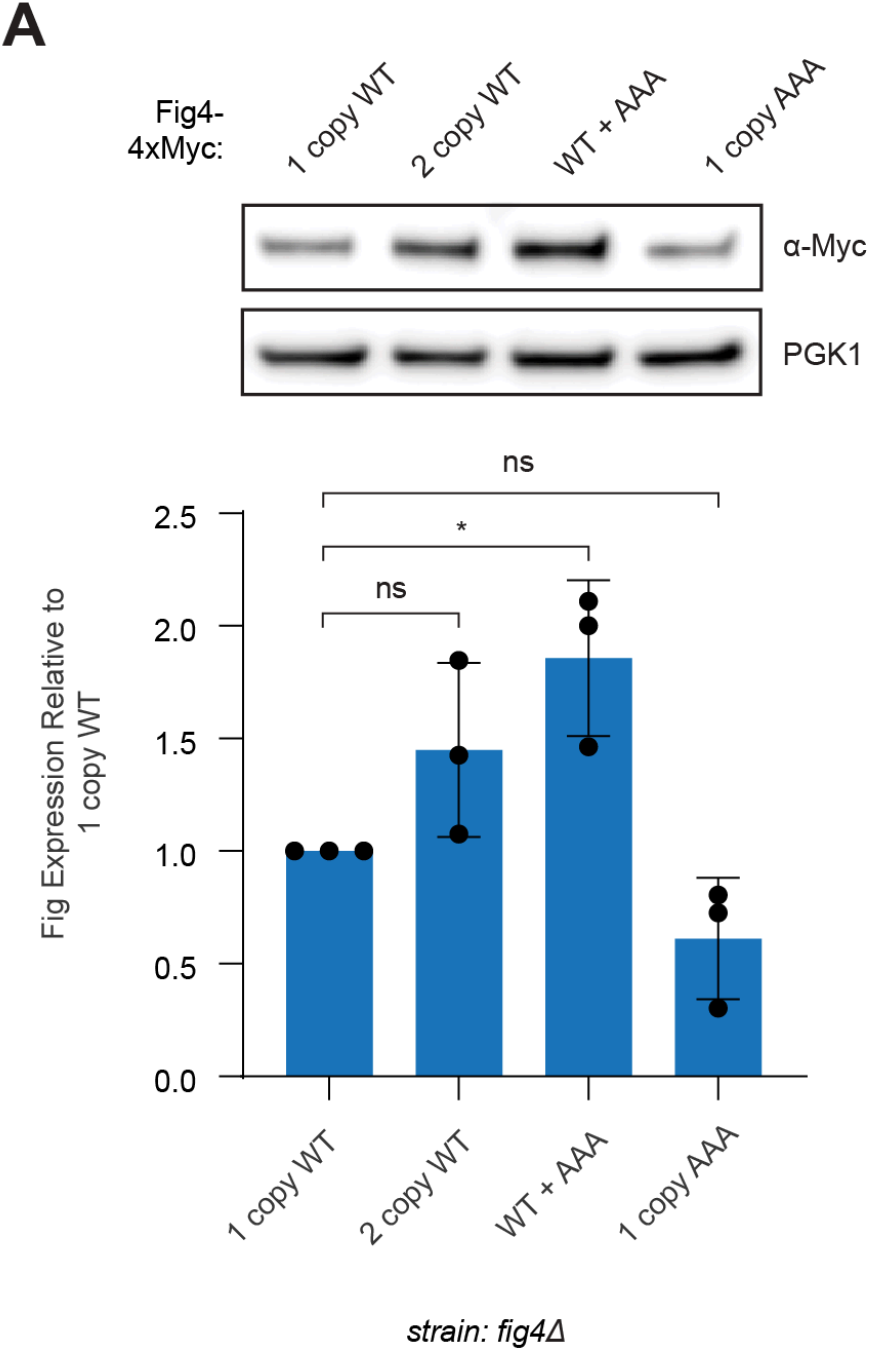
Relative levels of Fig4 in cells expressing one or two copies of Fig4. Western blot of proteins from denaturing lysis of a fig4Δ strain co-transformed with centromeric plasmids pRS415 (+pLEU2) and pRS413 (+pHIS3) expressing the indicated Fig4 4x-Myc-tagged variants or no Fig4 (vector). Plasmid based Fig4 was expressed with native promoters and 5′ and 3′ UTRs. Bar graph shows quantification of Fig4 band intensities in lysates relative to 1 copy of Fig4 wild-type. Error bars represent standard deviation (ns not significant, *p < 0.05 by two-tailed t-test).

**Figure S2.**
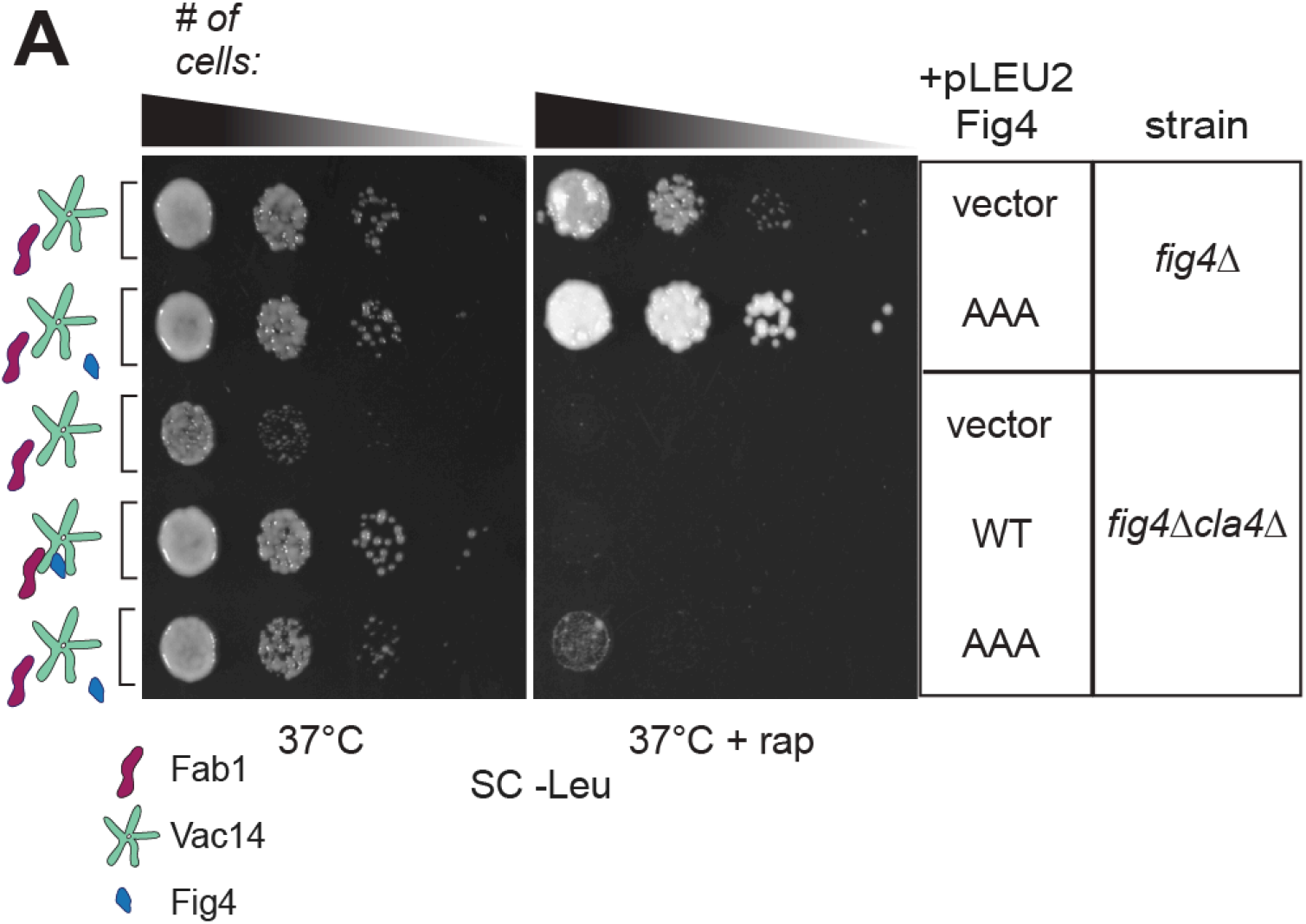
Rapamycin tolerance conferred by Fig4-AAA relative to Fig4-WT and no Fig4 persists in the absence of the Ste20 homolog Cla4. fig4Δ and fig4Δcla4Δ strains were transformed with pRS415 (+pLEU2) plasmids expressing the indicated Fig4 variants or no Fig4 (vector) as indicated. Plasmid based Fig4 was expressed with native promoters and 5′ and 3′ UTRs. Cells were spotted on SC-Leu agar plates in a 10-fold dilution series with or without 10 nM rapamycin and incubated at 37°C.

